# Nested tandem duplications of the gene *Melanoma antigen recognized by T-cells* (*Mlana*) underlie the sexual dimorphism locus in domestic pigeons

**DOI:** 10.1101/754986

**Authors:** Shreyas Krishnan

**Author notes:** Corresponding author: Shreyas Krishnan. Curent address: 2633 McKinney Ave, #130-352, Dallas, TX 75204.

## Abstract

Birds have classic examples of exaggerated sexually dimorphic traits, including colors. Sexes among the wild rock dove and its derived domestic breeds, however, are quite indistinguishable and sex can only be ascertained through genotyping or egg laying and successful hatching of eggs. Yet, the pigeon fancy has discovered sexually dimorphic traits and harnessed some of these traits in some auto-sexing breeds. Early genetics pioneers characterized the sex-linked *Stipper* locus and showed it to be linked to the pigeon Z linked *B-locus* (*Tyrp1*). The alleles of the *Stipper* locus have variable dominance relative to wild-type (more severe alleles are incompletely dominant, less severe alleles are reportedly fully dominant), characterized by a continuum of lightening in homozygotes and increased variegation in heterozygotes corresponding with severity. We leveraged this positional information and population structure among breeds in a candidate gene approach to identify the genetic mechanism of de novo pigeon sex dichromatism. A large tandem duplication (77 kb) centered on the gene *Melanoma antigen recognized by T cells* (*Mlana*) is completely associated with alleles of the *Stipper* locus. Copy number of the 77 kb genetic lesion was not correlated with allele severity suggesting that other mechanisms, including epigenetic regulation could underlie both allele severity and degree of variegation.

## Introduction

Sexual dimorphism is known from theoretical and empirical work to result from antagonistic natural selection or sexual selection acting on autosomal or sex-linked loci. Sexual dichromatism, especially is exaggerated to a greater extent among birds than among mammals, with cocks often being the targets for runaway sexual selection. Theory predicts that dichromatism is easier to evolve when a locus under selection is sex-linked by virtue of dosage effects in each sex (Rice 1984; Albert & Otto 2005), especially given the incomplete or lack of dosage compensation reported among birds (Teranishi et al. 2001; Shang et al. 2010; Caetano et al. 2014; Graves 2014).

In domestic pigeons, an otherwise uniform blue-gray species, classical genetic work has identified two pairs of sex-linked (Z-linked) color loci: *Stipper* and *B-locus*, and *dilution* and *reduced* (Cole & Kelley 1919; Hawkins 1931; Hollander & Cole 1940). *Stipper* (*St*) and *B-locus* (*b*) are closely linked (~3 cM), and approximately 45 cM from the second pair, *dilution* and *reduced* that are ~5 cM apart (Hollander, unpublished). Both *Stipper* and *B-locus* exhibit sexual dimorphism in degree of plumage variegation, and alleles of the former (e.g. *Faded*) have been fixed in some auto-sexing breeds (fig. 1). We recently showed two Z-linked genes, *Slc45a2* and *Tyrp1* map to the *dilution* locus and *B-locus* respectively (Domyan et al., 2014).

**FIG. 1.**
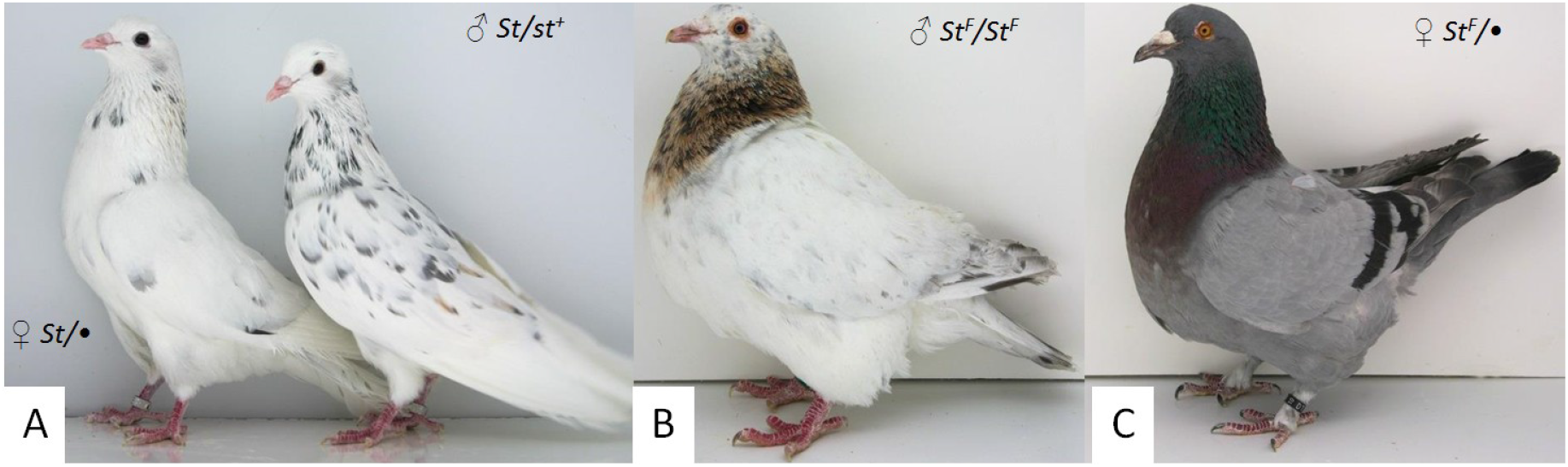
Sexual dimorphism and Variegation at the *Stipper* locus. **A**: Siblings from a nuclear family show dimorphism in degree of variegation. *Almond* individuals fledge white, but variegation and pigmentation increase with age; *Almond* females (left, 5 month old) develop less variegation than heterozygous males (right, 3 month old). The breeder’s classic *Almond* (not shown) in contrast to unrefined *Almond*s (shown here) combine mutant alleles in heterozygosity at several autosomal and sex-linked pigmentation loci that enhance the richness of the variegation. **B and C**: Texan pioneer is a squabbing breed (produced for meat), in which a *Faded* allele of a single origin has been fixed. **B**: a male blue Texan pioneer (wild-type/blue-gray at the *B-locus*) appears nearly white (*St^F^/St^F^*) in contrast to a female; **C**: a female blue Texan pioneer appears wild-type (*St^F^/•*).

In this study our goal was to map the *Stipper* locus, which is characterized by more extreme sexual dichromatism than the *B-locus*, including male-specific lethality for some of the more severe alleles in the homozygous state. Mapping the molecular underpinnings of the *Stipper* locus will provide a rare opportunity to examine the evolution of sexual dichromatism at its earliest stages. This locus is highly mutable in the germline, reportedly spontaneously giving off less severe alleles from the most severe allele (*Stipper/Almond*) in 1/100 informative meioses, and the *Faded* allele alone is reported to have arisen multiple times in this manner. The complexities of this locus are that almost all reports subsequent to Hollander’s work come from fancier-geneticists, and distinct phenotypic manifestations due to dosage, allele severity, somatic instability (variegation), and reported new alleles have likely arisen in previously unexamined genetic contexts (breeds previously not known for *Stipper* alleles). The *Stipper* locus has seven recognized alleles of decreasing severity and dominance: *Almond* or *Stipper* (*St*), *Sandy* (*S^Sa^*), *Qualmond* (*St^Q^*), *Hickory* (*St^H^*), *Faded* (*St^F^*), *Chalky* (*St^Ch^*), *Frosty* (*St^Fy^*), with wild-type (*st^+^*) at the bottom of the allelic series (Peter 2015). In homozygosity the phenotypes are reported to form a near-continuum of plumage lightening from blue-gray in wild-type to white in the *Almond* allele. *Almond* homozygotes, are reported to hatch out naked (without neonatal down), with defective eyes, and fledge white. These fledglings are not known to make it to sexual maturity. *Sandy* homozygotes unlike *Almonds* are reported to have mild eye defects, and are not homozygous lethal. In heterozygosity too phenotype severity is reported to correspond with dominance hierarchy, such that degree of variegation is greatest in *Almond*, and degree of variegation is greater in cocks than in hens. Variegation is least in homozygotes. Finally, each fleck of variegation increases in size and distribution with successive molts (fig. 1).

The *Almond* (*St*) allele in heterozygosity is prized among breeders for the rich colors produced in the flecks, particularly in combination with heterozygosity at a number of autosomal pigmentation traits and the linked *B-locus*. The pigmentation in the variegating flecks is influenced by the genotype and gametic phase at the neighboring *B-locus,* with distinct cis- vs. trans- allelic interactions in males doubly heterozygous at *Stipper* and *B-locus* (Hollander 1982). For example: in *Almond, B-locus* double heterozygotes, the flecks bear the color encoded by the dominant *B-locus* allele only when that allele is in trans- (out of phase) with the *Almond* (*St*) allele. When this phase relationship is reversed, the flecks may bear colors encoded by either *B-locus* allele.

The *Faded* (*St^F^*) allele has been harnessed by pigeon squab breeders for auto-sexing purposes in a small number of utility breeds, including the Texan pioneer (fig. 1 B & C) (Hollander 1982; Hollander 1938; Hollander & Cole 1940). This breed is unique in that the date and origin of the *Faded* allele in the breed is known (new breed admission 1962 NPA).

The large number of alleles, continuum of phenotypes, sexually dimorphic variegation and plumage lightening, and distinct cis- vs. trans- interaction with the *B-locus* are suggestive of instability in the vicinity of *B-locus*/*Tyrp1* that suppresses pigment synthesis or melanocyte differentiation. Several examples of continuous traits and variegation have been characterized in other species with multitude of mechanisms (Lyon 1962; Argeson et al. 1996; Kijas et al. 2001; Clark et al. 2006; Girton & Johansen 2008; Choate et al. 2010; Walbot 2013). As there are several models that might account for various aspects of these dimorphic variegation, identifying the molecular underpinning is essential to understanding these processes.

## Results

The Z-linked *Stipper* locus is reportedly 3 cM from *B-locus* (*Tyrp1*) and the chicken Z centromere is approximately 10 Mb from chicken *Tyrp1.* BLAST alignment of the pigeon Z centromeric PR1 repeat and the chicken CENP-A, Z centromeric repeat sequences map across pigeon scaffold 6 approximately 3 Mb upstream from *Tyrp1* (scaffold 6 – 4.8 Mb; *Tyrp1* – 204968-215057) (Solovei et al. 1996; Shang et al. 2010). A CR-1 non-LTR transposable element was discovered interrupting the centromere repeat alignment when BLASTing a 2 Kb sequence of pigeon genomic scaffold 6, centered on the PR1 repeat against the pigeon genome. The *B-locus*, *Ash-red* haplotype spans the proximal portion of scaffold 6 and includes centromeric repeats at its distal 5’ end [Domyan et al., (2014) refined this to a 20 kb haplotype; (Solovei et al. 1996; Shang et al. 2010)].

Chromosome walking outward of *Tyrp1* along flanking scaffolds in Texan pioneer pigeons (*St^F^*/*St^F^*) formally excluded this gene (fig. 2). Texan pioneers from a single loft in all three *B- locus* alleles/haplotypes were tested, revealing shorter than expected haplotypes than if this gene was casual. On the 5’ of *Tyrp1*, the tested markers were not polymorphic at the end of scaffold 6, suggestive of the predicted long runs of homozygosity expected around the *Faded* (*St^F^*) haplotype; candidate gene *Melanoma antigen recognized by T cells 1* (*Mlana*) resides at 3.17 Mb on scaffold 6 and *Adaptor-related protein complex 3, beta-1 subunit* (*Ap3b*, scaffold 215) is located on a flanking scaffold (fig. 2). Sequencing exons of candidate gene *Mlana* during an earlier study among unrelated individuals of several breeds showed lack of polymorphism in a panel that included *Stipper* phenotypes.

**FIG. 2.**
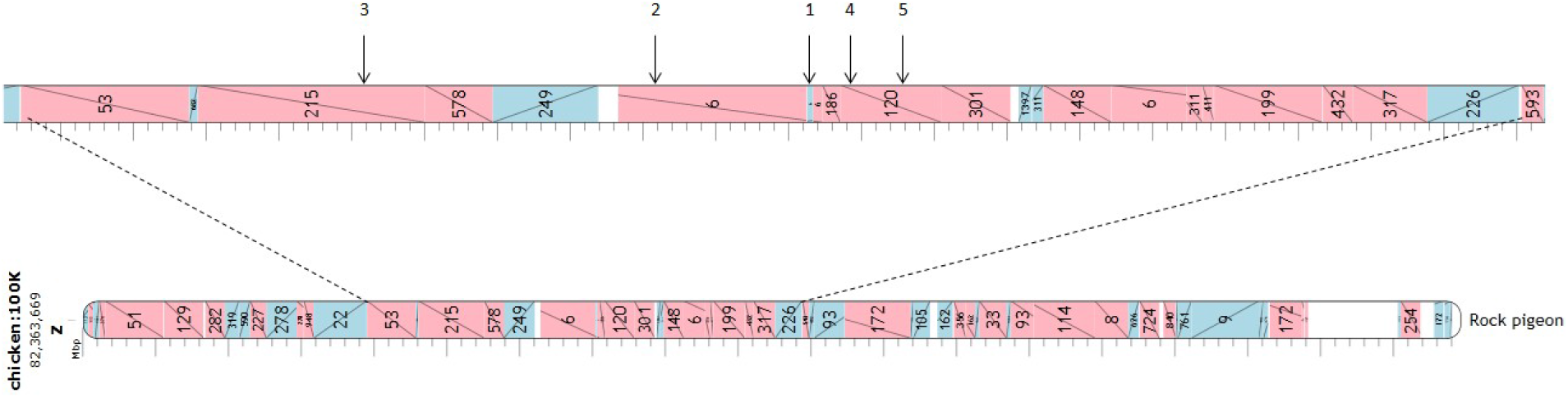
Pigeon scaffolds aligned to Chicken Z chromosome (Ggal 4.0), indicating scaffold orientation and relative scale using the interactive Evolution highway chromosome browser (v1.0.6002, http://evolutionhighway.ncsa.uiuc.edu/). Numbered arrows in the top panel represent relative positions of candidate genes for the *Stipper* locus: 1) *Tyrp1*, 2) *Mlana*, 3) *Ap3b1*, 4) *Nfib*, 5) *Ledgf*.

We investigated scaffolds in this region of all resequenced pigeon genomes that bear candidate genes for structural polymorphism by piling up the regional depth over respective library genome wide averaged read depth. Two putative, large, nested, copy number variants (CNV) 5’ of *Tyrp1*, both centered on candidate gene *Mlana* (scaffold 6) were discovered in several libraries. The smaller 25 kb tandem duplication, spanning only *Mlana* occurred in ten libraries, while the larger 77 kb tandem duplication occurred in five of those ten libraries (fig. 3). The 77 kb duplication appears to have occurred on the 25 kb duplication haplotype and does not appear from the WGS data to crossover and segregate away. It is thus predicted that a Z chromosome bearing the 25 kb duplication has two copies of *Mlana* and a chromosome bearing the 77 kb duplication is expected to have four copies of *Mlana*. None of the breed standards for libraries having either CNV were suggestive of selection for specific *Stipper* alleles. However, the possibility that some breeds are fixed for traits that may actually be unrecognized alleles of *Stipper* is considered. Given the high frequency of the 25 kb CNV, we chose to empirically validate and test the 77 kb duplication with a panel of *Stipper* allele individuals. The distinguishing feature among each haplotype and wild-type are the breakpoint spanning sequences. Reads for the region flanking the boundaries of the putative CNVs were manually aligned to identify the breakpoint spanning sequences across which allele specific amplicons were designed. A large CT rich repeat (TTTCCCTTTTCCTTCCTTTTCCCCC)_~25_ flanking the breakpoint and the duplication specific primer of the 77 kb CNV allowed only a presence/absence assay. The 77 kb duplication was empirically verified in a panel comprising unrelated *Almond* cocks and individuals of *Sandy, Qualmond, Faded, Chalky,* and *Frosty* alleles. Genetic association was tested in an expanded panel comprising wild-type hens and cocks and *Stipper* alleles from unique lineages. The breakpoint spanning amplicon had PCR amplification in all of the *Stipper* individuals, and failed to amplify in any of the wild-type individuals (Fisher’s exact test: p = 3.56 × 10^−15^; fig. 4).

**FIG. 3.**
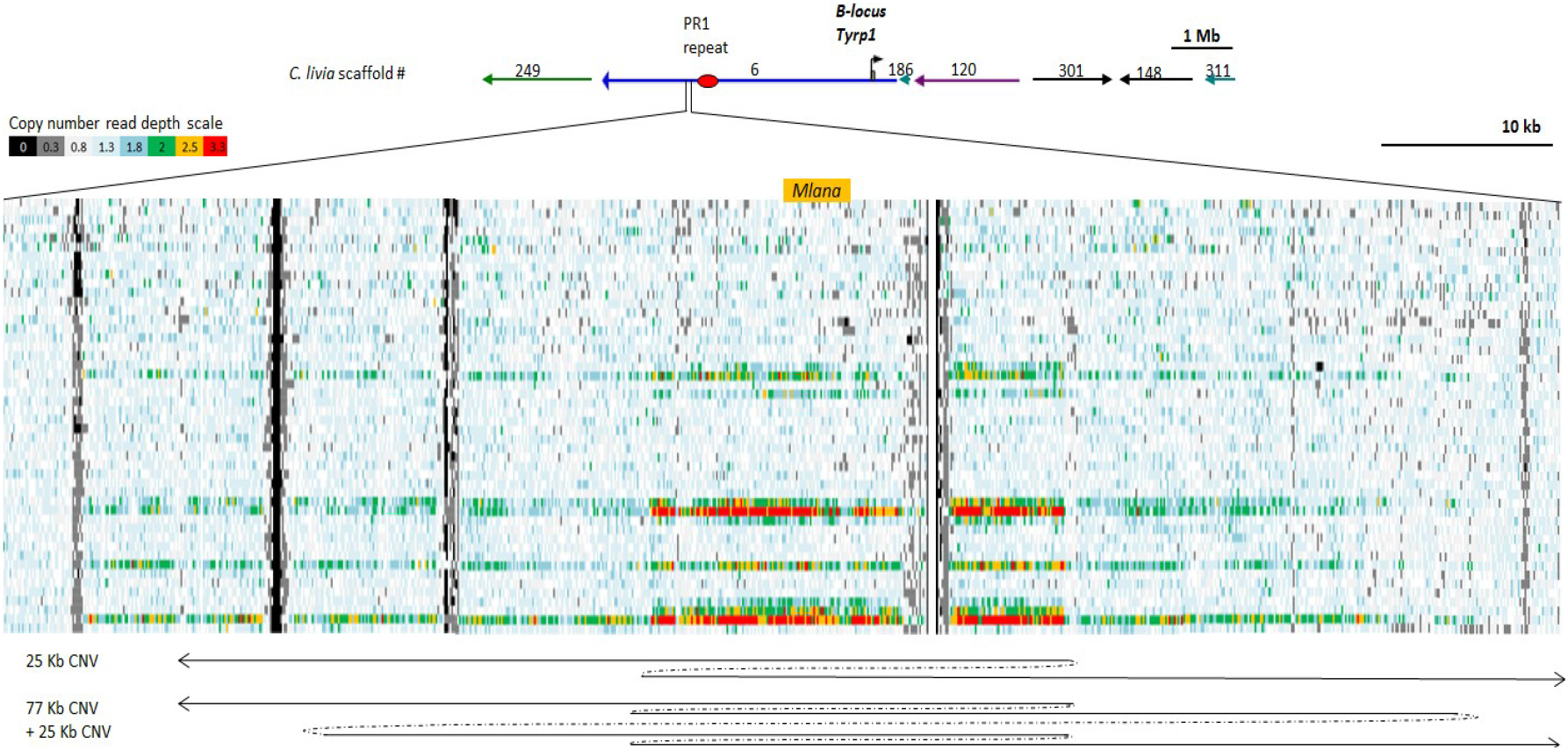
Elevated read depth reveals nested serial amplifications of *Mlana*. Scaffolds in the vicinity of *Tyrp1* (a linked locus) were examined for variation in read depth: average depth in 100 bp sliding windows normalized to library-specific genome-wide averages were plotted. The above panel presents a region of scaffold 6 (3127701-3230001) that also bears *Tyrp1* [Scaffold 6– 4.8 Mb, *Mlana* coordinates – 3,173,417-3,178,870, centromeric repeats on scaffold6 are labelled PR1 repeat]. Elevated read depth spanning 25 kb centered over *Mlana* (scaffold 6) was observed in ten out of 40 WGS libraries (Berlin long face tumbler, SRR516979; Birmingham roller, SRR516980; Chinese owl, SRR516982; African owl, SRR516994; Ice pigeon, SRR516995; Frillback, SRR516996; Lahore, SRR517002; Archangel, SRR517006; Cumulet, SRR517007; Egyptian swift, SRR517008). Five of these ten libraries had further increase in read depth also centered on *Mlana*, but spanning 77 kb (Birmingham roller, SRR516980; African owl, SRR516994; Ice pigeon, SRR516995; Lahore, SRR517002; Egyptian swift, SRR517008). Order of libraries and breed names from top to bottom row is indicated in same order in methods section.

**FIG. 4.**
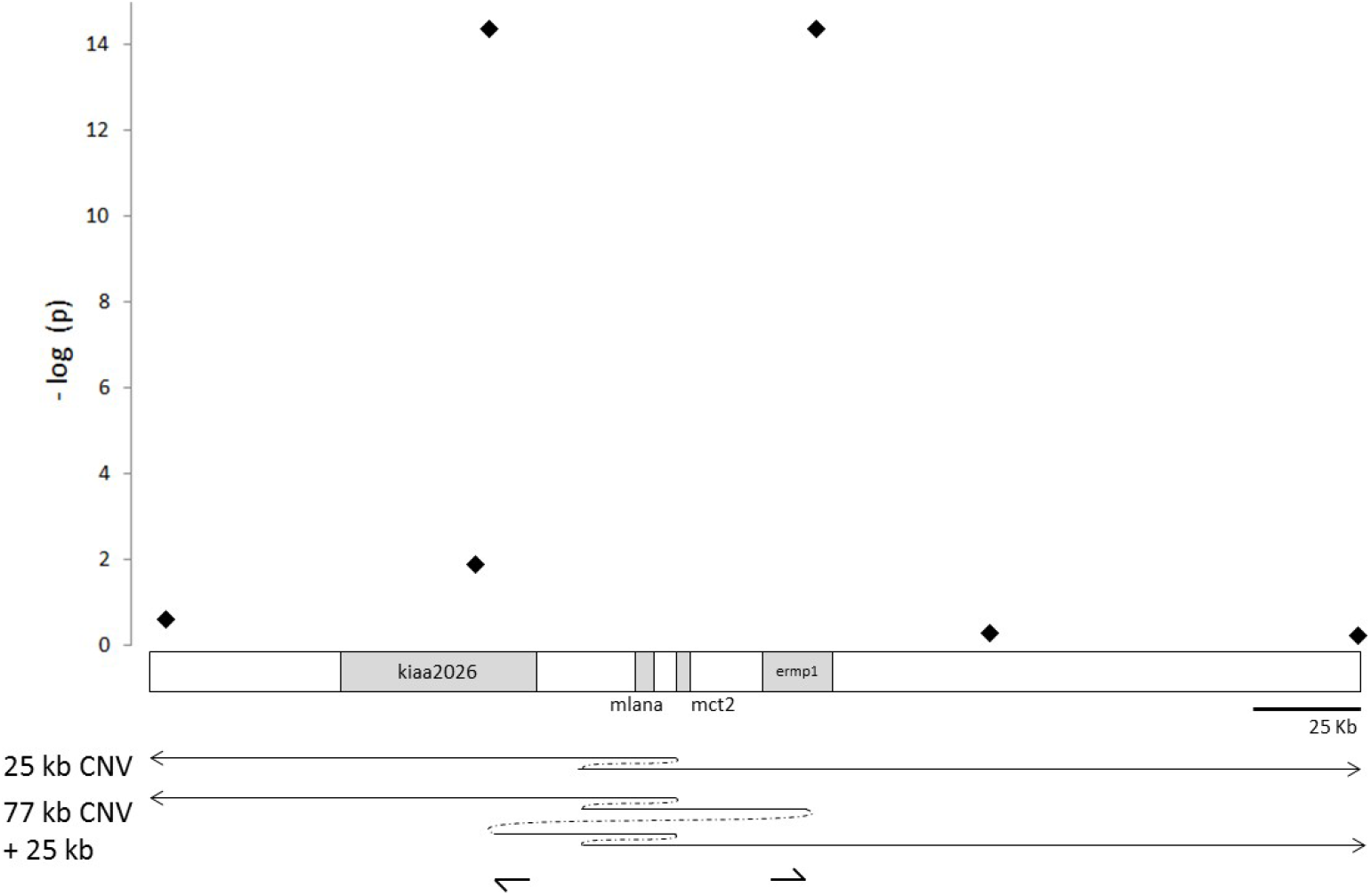
Bounding the associated critical interval for *Stipper*. The significance of the association of markers in the vicinity of *Mlana* with *Stipper* is plotted as −log (p). The two high-significance points represent the breakpoint-spanning amplicon marker (primers indicated by arrows). Schematic under the graph of significance illustrates the structure of the 25 kb and nested 77 kb + 25 kb CNVs.

The 77 kb duplication spans only *Mlana* and *Monocarboxylate transporter 2* (*Mct2*) in their entirety, as well as portions of two additional genes. To bound the critical interval around the 77 kb duplication for association with *Stipper*, indel markers were genotyped at intervals walking away from the duplication breakpoints in *Stipper* mutant and wild-type individuals. Recombinant haplotypes were identified 3 kb from the left flank (Fisher’s exact test: p = 0.01) and 44 kb from the right flank of the 77 kb duplication (Fisher’s exact test: p = 0.08; fig. 4).

In a breakpoint spanning assay to test the association of the smaller 25 kb duplication in a subset of the previous association panel, this CNV was shown to be weakly associated with alleles of the *Stipper* locus (Fisher’s exact test: p = 0.007).

### Copy-number evaluation

To test the correlation of copy number and *Stipper* allele severity a wild-type hen was ascertained by testing the absence of both CNVs (25 kb and 77 kb). An *Almond* hen was used as positive control (amplification fold change relative to wild-type control: 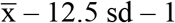, n – 4 technical replicates) (fig 5). Validation of the copy-number assay was based on the premise that the *Faded* allele of Texan pioneer pigeons will have a narrow range of variation due to their known single origin. Using the method of Livak and Schmittgen (2001), Texan pioneer hens (n = 2) tested had seven-fold amplification of *Mlana* in contrast to 13-fold and 16-fold amplification in two unrelated *Almond* hens, relative to an ascertained wild-type hen. Copy-number (fold change) of *Mlana* per chromosome ranged from three to 18 in *Almonds* (*Almond* cock: 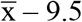, sd – 6.2, n – 4; *Almond* hen: 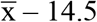, sd – 2.1, n – 2) and four to seven in *Faded* (*Faded*: 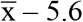, sd – 1.6, n – 4) (assuming their wild-type chromosome bears a single copy *Mlana* in heterozygotes, and both chromosomes bear equal numbers of *Mlana* in homozygotes). Copy-number of *Mlana* among unrelated cocks homozygous for *Frosty, Chalky, Faded,* and *Sandy* increased in an apparent linear fashion, however, only a single specimen was available for these alleles. Within a nuclear family sired by an *Almond* cock copy-number ranged from six to 17 per chromosome (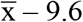, sd – 5.02, n – 5).

**FIG. 5.**
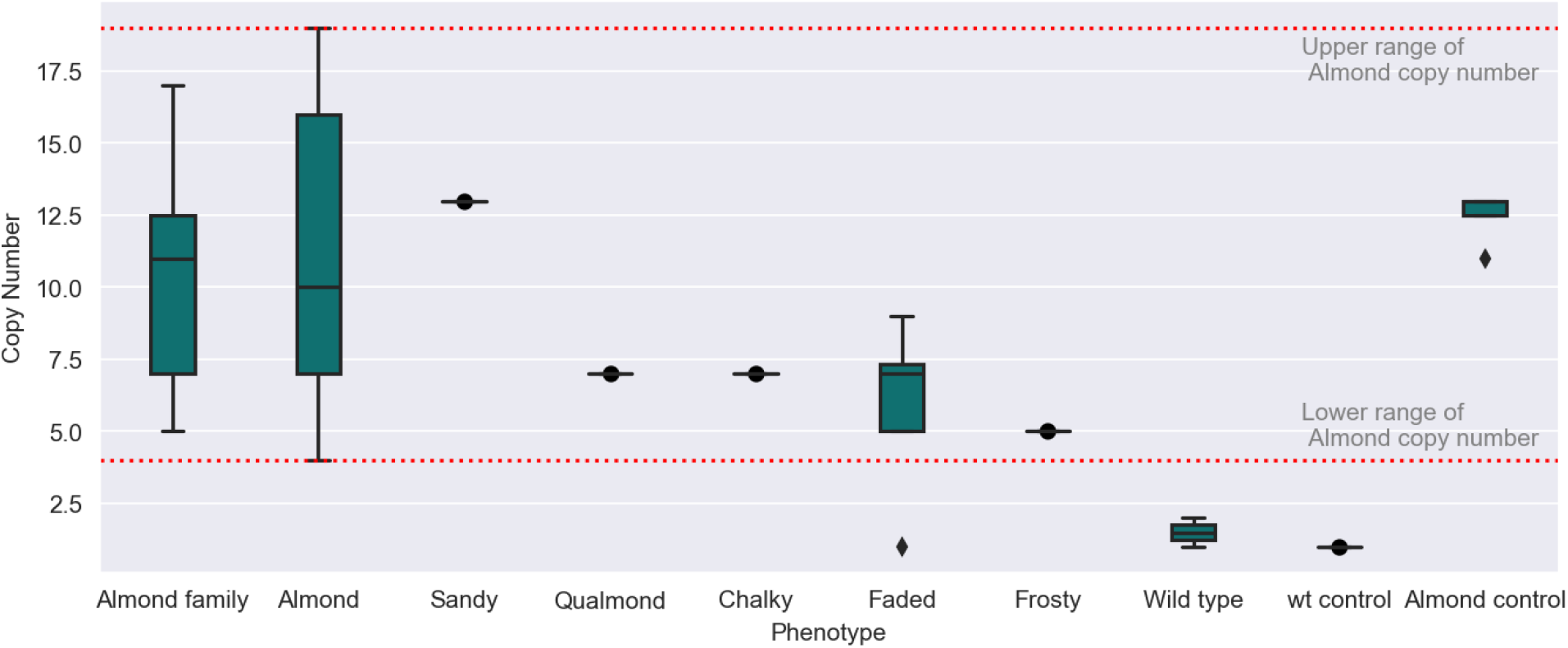
*MLANA* haploid copy number of *Stipper* locus phenotypes ascertained by SK or JWF, and estimated by the 2^−ΔΔCT^ method. Phenotypes are ordered in decreasing order of severity as reported by fanciers. *Almond* control (positive control) was a hen hatched in our lab from the *Almond* nuclear family (also shown). Data for *Almond* phenotypes includes unrelated individuals and only the sire of the *Almond* family.

## Discussion

De novo origins of sexual dimorphism are without doubt of great interest to biologists, especially when we can observe and test the molecular underpinnings. Pigeons provide a unique opportunity to understand the underpinnings of sex-linked dimorphism. These dimorphic pigmentation traits were observed by early pioneers including Darwin, and are reported to have been harnessed even in ancient breeds like the Reehani dewlap to discriminate sexes (Darwin 1868; Levi 1963). Four sex-linked pigmentation traits were recognized by early geneticists and the sexually dimorphic expression was recognized for two of the four loci. The positional information of the four sex-linked loci together with advances in our understanding of the melanogenesis pathway and availability of the pigeon genome data and well annotated chicken genome limited the genome-wide pool of candidate genes on the Z chromosome.

Given the reported 3 cM distance of the *Stipper* locus from *B-locus* and similarities between the *Ash-red* phenotype at *B-locus* and alleles of *Stipper*, we have shown here that a 77 kb tandem duplication centered on *Mlana* is in complete association with alleles of the *Stipper* locus (p = 3.56 × 10^−15^). The 25 kb tandem duplication spans only *Mlana*, whereas the 77 kb tandem duplication also spans *Monocarboxylate transporter 2* (*Mct2* or *Slc16a7*) and portions of KIAA2026 and *Endoplasmic reticulum metallopeptidase 1*. MCT proteins are a poorly understood family of solute carriers that transport monocarboxylates such as pyruvate, lactate, and ketones across cell membranes for carbohydrate, lipid, and amino acid metabolism (Halestrap & Wilson 2012). *Mct2* has different expression profiles in the few species in which it has been studied and is not reported to be expressed in melanocytes (Halestrap & Wilson 2012). *Mlana*, the only pigmentation candidate in the region is a melanoma, melanocyte and retinal pigment epithelial cell marker (Coulie et al. 1994; Aydin & Beermann 2009; Aydin et al. 2012). The MLANA protein is reported to play a key role in maturation of Premelanosome protein (PMEL), a melanosomal matrix protein, upon which the melanin pigments are deposited (Hoashi et al. 2005).

The 77 kb genetic lesion with *Mlana* at its center is the seed for the allelic continuum at *Stipper* locus. While there is an indication of a linear relationship between copy-number and allele severity among homozygotes of *Stipper* alleles, lack of a clear copy-number correlation with *Almond* allele severity may be a consequence of phenotype ascertainment bias or genetic heterogeneity and other models that are discussed below.

Intriguingly, the *ice* pigeon library (SRR516995), which is known to be fixed for the *ice* phenotype has both the 77 kb and 25 kb duplications. The breed is additionally fixed for wild-type (blue) at the *B-locus.* Considering this breed is not known to exhibit variegation or sexual dimorphism and has no reports of sub-lethal effects or reduced fertility, it is conceivable that *ice* could be an allele of the *Stipper* locus, but lower than *Faded* in dominance order and severity.

### *Models to explain attributes of* Stipper *mutants*

Several non-exclusive models account for multiple features of this locus. Simple sex-linked dosage effects could manifest in different character states in each sex and genotype because in birds, unlike in mammals, global silencing of one sex chromosome in the homogametic sex is not known to occur (Lyon 1962; Teranishi et al. 2001). The extent and mechanisms of avian dosage compensation are still unclear, and are areas of active research, but recent results in chickens indicate that a chromosomal interval flanking *Dmrt1* (candidate avian sex determination locus) is hyper-methylated in males and hypo-methylated in females by the long non-coding RNA of the *Male hyper-methylation* locus (*MHM*) (Teranishi et al. 2001; Shang et al. 2010; Caetano et al. 2014; Graves 2014). This mode of regional dosage compensation found in chickens does not appear to be found in all birds, as demonstrated by the lack of the *MHM* locus in zebra finches (Itoh et al. 2010).

The continuous nature of the plumage lightening phenotypes of various *Stipper* alleles, however, is suggestive of dosage modulation due to copy number variation. The bioinformatic data leads one to expect a chromosome bearing a single 77 kb duplication to have four copies of *Mlana*. The copy number assays, however, reveal a wide range of variation among alleles of *Stipper*, especially among unrelated *Almond,* and even among related *Almond* individuals.

The variegation in the *Stipper* phenotypes is reminiscent of position effect variegation (PEV) in *Drosophila* and the mouse *Agouti hypervariable* allele (Talbert & Henikoff 2006; Slotkin & Martienssen 2007), in both of which a genetic lesion triggers epigenetic modifications in the soma that restores the underlying phenotype. Similarly, if epigenetic modifications of the 77 kb genetic lesion are inherited stably in the germ-line, then the allelic continuum and heritability of the phenotype could be easily explained (fig. 1A: copy number of the hen/positive control = 13, and cock = 4). In the melanocytes, variation in copy number together with its epigenetic modifications during feather growth could result in variegation and explain the allelic continuum and cis−/trans - effects.

The long stretch of tandemly repeated homologous sequence is also ripe for illegitimate recombination, and can also result in expansion or contraction of the motif in the germline, similar to well-known examples in dogs and humans (Fondon & Garner 2004; Choate et al. 2010; Gemayel et al. 2010). Such somatic recombination models could adequately account for the *cis*- vs. *trans*- allelic interactions between the *Stipper* and *B-locus.* A simple model of repeat expansion and contraction can however, entail a runaway process that can cause breakage of the chromosome that does not occur here (based on markers examined near the chromosome tails). A “fragile Z” model might also explain variegation, but may not be adequate on its own to explain the *cis*- vs. *trans*- allelic interaction between alleles of the *Stipper* and *B-locus* in the soma unless *Tyrp1* is centromeric to *Mlana*. It is conceivable that a combination of CNV expansion and somatic recombination can underlie the basis of allelic continua, de novo origins of alleles in pedigrees, and variegation.

An alternate model considers the requirement of the MLANA protein to participate in the maturation of the PMEL. In dogs, a SINE insertion in the terminal intron of *Pmel* is associated with the merle trait, which is a patchwork of wild-type and depigmented patches (Clark et al. 2006). Similarly loss of function mutations of *Pmel* in chicken, horse, and cattle result in varying degrees of hypopigmentation (Kerje et al. 2004; Brunberg et al. 2006). If the native state of the 77 kb *Mlana* tandem duplications in the germline is epigenetically silenced, then copy-number of *Mlana* may not be crucial to the phenotype as the data seem to suggest. In this scenario, silencing of *Mlana* can have downstream effects on PMEL and melanosome morphology, thus resulting in the bleached phenotype, such that loss of silencing in the germline and soma results in de novo lower alleles and variegation (Aydin et al. 2012).

One intriguing possibility is raised by the fact that MLANA is a melanoma specific antigen recognized by cytotoxic T lymphocytes (CTL). It is thus conceivable that over-expression of *Mlana* triggers an auto-immune response resulting in immune-mediated depletion of melanocytes. Sufficient exposure to CTLs, which is known to select for cells that do not express the antigens (Jäger et al. 1996), may relieve melanocyte depletion and restore melanogenesis in the pigmented flecks of feathers.

## Methods

Foundation stock was obtained from hobbyists to screen for marker-phenotype association; in addition crosses were established between diverse breeds in which to test co-segregation of markers and color phenotypes. Blood was drawn from the brachial or axillary vein and stored in ice in EDTA vacutainers, or blood was drawn into heparinized capillary tubes and plunged into 50% ethanol in 1.5 mL centrifuge tubes. All animals were photographed in a customized light box standardized for dimensions, illumination, focal length and field of view using a Nikon D5000 DSLR camera. Detailed interviews were conducted with breeders for each sample to ascertain the phenotypes, genotypes, and to assess confidence. Pedigree information including phenotypes/genotypes of first degree relatives, and information on size and diversity of loft, and breeding goals of the breeder were used to vet and inform decisions on sample ascertainment and confidence. Knowledge of the Z linkage group was used to target samples across breeds such that phase relationships could be deduced with the objective that mapping one locus will help lead to nearby loci. All experiments involving animals were conducted in accordance with Institutional Animal and Use Committee guidelines (protocol #s A09.009 & A14.009; PI: JWF).

In the chicken Z chromosome (Ggal 4.0) *Tyrp1* (the pigeon *B-locus*) lies 10 Mb from the metacentric centromere. Published pigeon and chicken centromeric repeat sequences from Genbank (Solovei et al. 1996; Shang et al. 2010) were BLASTed against the pigeon genome to determine the location of the pigeon Z centromeric repeats in relation to the pigeon *B-locus* (*Tyrp1*, scaffold 6). Gene order and orientation were screened by eye and gene order at scaffold termini were used to determine scaffold ordering by contrasting against the chicken and hoazin genomes, in addition to using the chromosome bowser (Evolution Highway v1.0.6002). Whole genome shotgun sequence (WGS) reads were extracted using Samtools and manually aligned to bridge across scaffold breakpoints in order to identify candidate polymorphisms. Candidate genes that account for different attributes of *Stipper* in this chromosomal interval include *Tyrp1* (pigeon *B-locus*) and *Melanoma A (Mlana*, melanoma antigen recognized by T cells) (Aydin & Beermann 2009; Aydin et al. 2012). Additionally, *Nuclear factor I B* (*NfIb*, melanocyte stem cell homeostasis), *Lens epithelium derived growth factor* (*Ledgf*, maintenance and survival of lens epithelial cells), *Adapter protein complex 3B subunit 1* (*Ap3b1*, adaptor protein complex*-Tyrp1* transporter) lie on flanking scaffolds and were considered (fig. 2; Nakamura et al. 2000; Pietro et al. 2006; Chang et al. 2013; Domyan et al. 2014).

To bound the chromosomal interval of the *Stipper* locus relative to *B-locus*, markers (SNP and indel) were selected from WGS data of Shapiro et al. (2013), and genotyped to exclude *Tyrp1* in association panels comprising closed sub-populations with confidence of few or single origins of the *Stipper* allele. To determine the direction in which *Stipper* lies, chromosome walking on either side of *Tyrp1* was performed in the Texan pioneer breed for homozygosity mapping among individuals of unknown relationships from a single loft. Since all three *B-locus* alleles occur in the breed, Texan pioneers are therefore expected to share the same, albeit large haplotype around *Faded*, but harbor variation at the *B-locus*.

### Detection of variants from whole genome shotgun sequence libraries

Available sequence reads from whole genome shotgun sequence libraries from 40 pigeons of diverse breeds/varieties (described in Shapiro *et al.* 2013) were obtained from the Sequence Read Archive (SRA, BioProject accession PRJNA167554) and used to identify sequence variation within Z chromosome scaffolds. The SRA format was converted into split FASTQ files for mapping using the fastq-dump tool included in the SRAtoolkit provided by NCBI (https://github.com/ncbi/sra-tools/wiki/Downloads). These FASTQ formatted reads were mapped to the Cliv1.0 pigeon reference genome using the Novoalign short-read mapping software, version 2.08.02 (Novocraft Technologies, Selangor, Malaysia) with default parameters. Single nucleotide variants and short insertions and deletions were identified using the mpileup feature of SAMtools (Li *et al.* 2009). Simple sequence repeat length variations were characterized using the methods of Fondon et al. (2012), for repeats identified by Tandem Repeat Finder v2.30 (Benson 1999) with parameters “2 5 5 80 10 14 5.”

Larger insertions, deletions, and other structural variation within 50 kilobases of *Mlana* were identified by anomalous inferred insert size, mate-pair orientation, or mapped read depth retrieved using the mpileup feature of SAMtools (Li *et al.* 2009). Read depth per base position in a 100 kb window centered on each candidate gene was plotted and examined to identify structural variation in the resequenced genome panel. A second approach plotted average read depth per 100 bp sliding window normalized over the genome wide average read depth for the respective library to examine gross structural variation that could be missed in the first approach. Phenotypes of the libraries-breeds used were not available, however, knowledge of breed standards and breeding practices allows us to determine which breeds segregate for *Stipper* phenotypes. Of the breeds listed none are fixed for known *Stipper* phenotypes. Order of libraries and *Columba livia* breed names used (also see Fig 3): Danish Tumbler (SRR511892, SRR511911, SRR511912, SRR511913, SRR511914, SRR511915); Indian Fantail1, SRR516967; American Fantail, SRR516968; *Columba rupestris* (SRR516969, SRR516970, SRR516971); Racing homer, SRR516972; Oriental frill, SRR516973; Laugher, SRR516974; Indian fantail, SRR516975; Feral1, SRR516976; English carrier, SRR516977; Scandaroon, SRR516978; Berlin long face tumbler, SRR516979; Birmingham roller, SRR516980; King, SRR516981; Chinese owl, SRR516982; Saxon monk, SRR516983; Syrian dewlap, SRR516984; Shakhsharli, SRR516985; Oriental roller, SRR516986; Carneau, SRR516987; English long face tumbler, SRR516988; English pouter, SRR516989; Jacobin, SRR516990; Mookie, SRR516991; Starling, SRR516992; Spanish barb, SRR516993; African owl, SRR516994; Ice pigeon, SRR516995; Frillback, SRR516996; Iranian tumbler, SRR516997; Marchenero pouter, SRR516998; Saxon pouter, SRR517000; English trumpeter, SRR517001; Lahore, SRR517002; Lebanon, SRR517003; Parlor roller, SRR517004; Racing homer, SRR517005; Archangel, SRR517006; Cumulet, SRR517007; Egyptian swift, SRR517008; Feral2, SRR517009. Finally, to bound the *Stipper* associated haplotype, markers were genotyped with the expectation that a recombinant haplotype will exclude flanking regions.

### Copy number assay

To empirically verify existence of copy number variants, primers were placed across breakpoint spanning sequences and tested in a discovery panel comprising various *Stipper* alleles. Allele specific tailed primers were designed for use in a three primer PCR assay, controlling for oligonucleotide sequence, amplicon sequence, genomic context, while preserving haplotype specific features and microhomology sequence between the allele specific primers.

Genotyping was performed using polymerase chain reaction (PCR) standardized for 10 μL volume reactions, 0.1 μL Taq DNA polymerase, 0.5 μL left and right primers (40 μM) respectively, 5 μL Failsafe premix (EpiCentre), 2.9 μL H_2_O, and 1 μL genomic DNA template. PCR cycling conditions were 96 °C – 3 min, (96 °C – 0.30 min, (65 – 55 °C) 0.30 min, 72 °C – 0.30 min) × 29, 72 °C – 6 min, hold at 4 °C, or touchdown protocol [96 °C – 3 min, {(96 °C – 0.30 min, 65-55 °C (−0.5 °C) – 0.30 min, 72 °C – 0.30 min) × 11}, {(96 °C – 0.30 min, 55 °C– 0.30 min, 72 °C – 0.30 min) × 30}] 72 °C – 6 min, hold at 4 °C].

#### Copy-number assessment

To evaluate copy-number correlation with allele severity, the Applied Biosystems ® SYBR green I chemistry (Power SYBR and Fast SYBR) were employed on an ABI7300 instrument. Quantitative PCR was performed using manufacturer’s protocol for 20 μL volume reactions, 10 μL SYBR, 0.2 μL left and right primers (20 μM) respectively, 7.6 μL H_2_O, and 2 μL genomic DNA template. Cycling conditions for Power SYBR and Fast SYBR on the ABI 7300 were 95 °C – 10 min, [(95 °C – 0.10 min, 60 °C – 0.30 min) × 40] and 95 °C – 0.23 min, [(95 °C – 0.10 min, 60 °C – 0.30 min) × 40] respectively. Relative quantification of copy number was assessed using the 2^−ΔΔ*C*T^ method, which contrasts PCR signal at the target or candidate region with an untreated control locus of known copy-number (Livak & Schmittgen 2001). ΔCT values for each sample were normalized against those of the wild-type hen to estimate fold change in amplification efficiency. Two amplicons of 90 bp length were designed, placing one amplicon in the heart of the candidate region (*Mlana* exon 4) and the second amplicon in a region on the Z chromosome that was ascertained bioinformatically to be invariant. To ensure wild-type genotype (single copy of *Mlana* per chromosome), phenotypically wild-type hens were genotyped for absence of the 25 kb and 77 kb duplication. An ascertained wild-type hen was used for further experiments. To ascertain the reliability of this method a panel comprising a wild-type hen, *Almond* hen (*St/•*) and two unrelated Texan pioneer hens (*St^F^/•*), was used with the expectation that the Texan pioneers will have comparable amplification fold change relative to wild-type and half that of *Almond*. Follow-up experiments were designed to evaluate relative copy numbers of various alleles of the *Stipper* locus in males (homozygotes and heterozygotes) and females. Finally, to independently ascertain the relationship of copy-number and *Stipper* phenotype all *Almond* individuals from a nuclear family sired by an *Almond* cock were assayed.

## Acknowledgments

This project was part of SK’s dissertation work (Dec 2016) and was supported through start-up funds to John W Fondon III (JWF) from the University of Texas at Arlington. Molecular work was done by SK and bioinformatic work was done by SK guided by JWF. All experiments involving animals were conducted in accordance with Institutional Animal and Use Committee guidelines (protocol #s A09.009 & A14.009; PI: JWF). Sample collection and animal facility management was possible with the help of several interns and the UTA animal facility staff. Several breeders including Richard Cryberg, Kerry Hendricks, and Dan Smith contributed samples and their wisdom from breeding and studying the loci mentioned here.

## Appendix

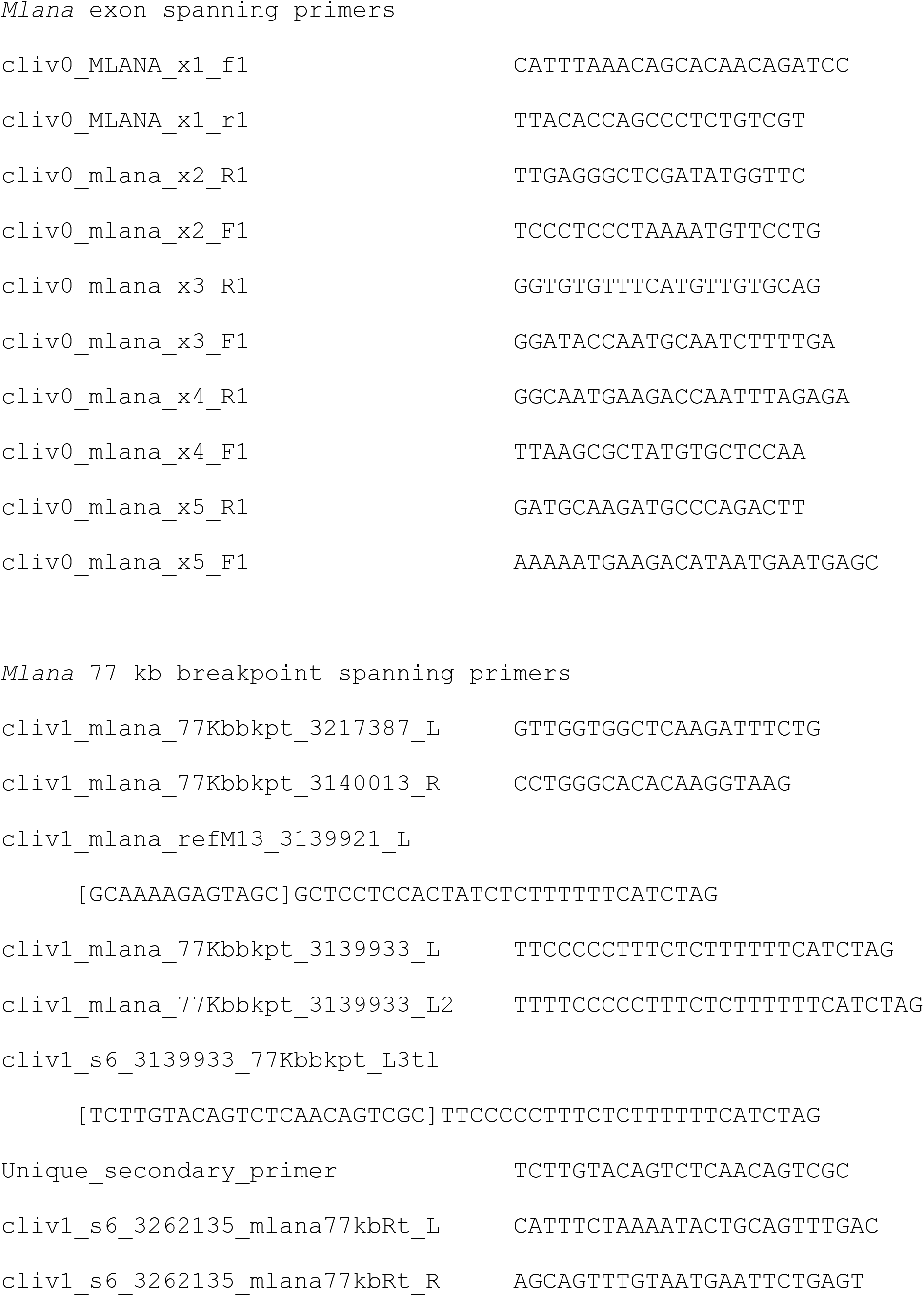
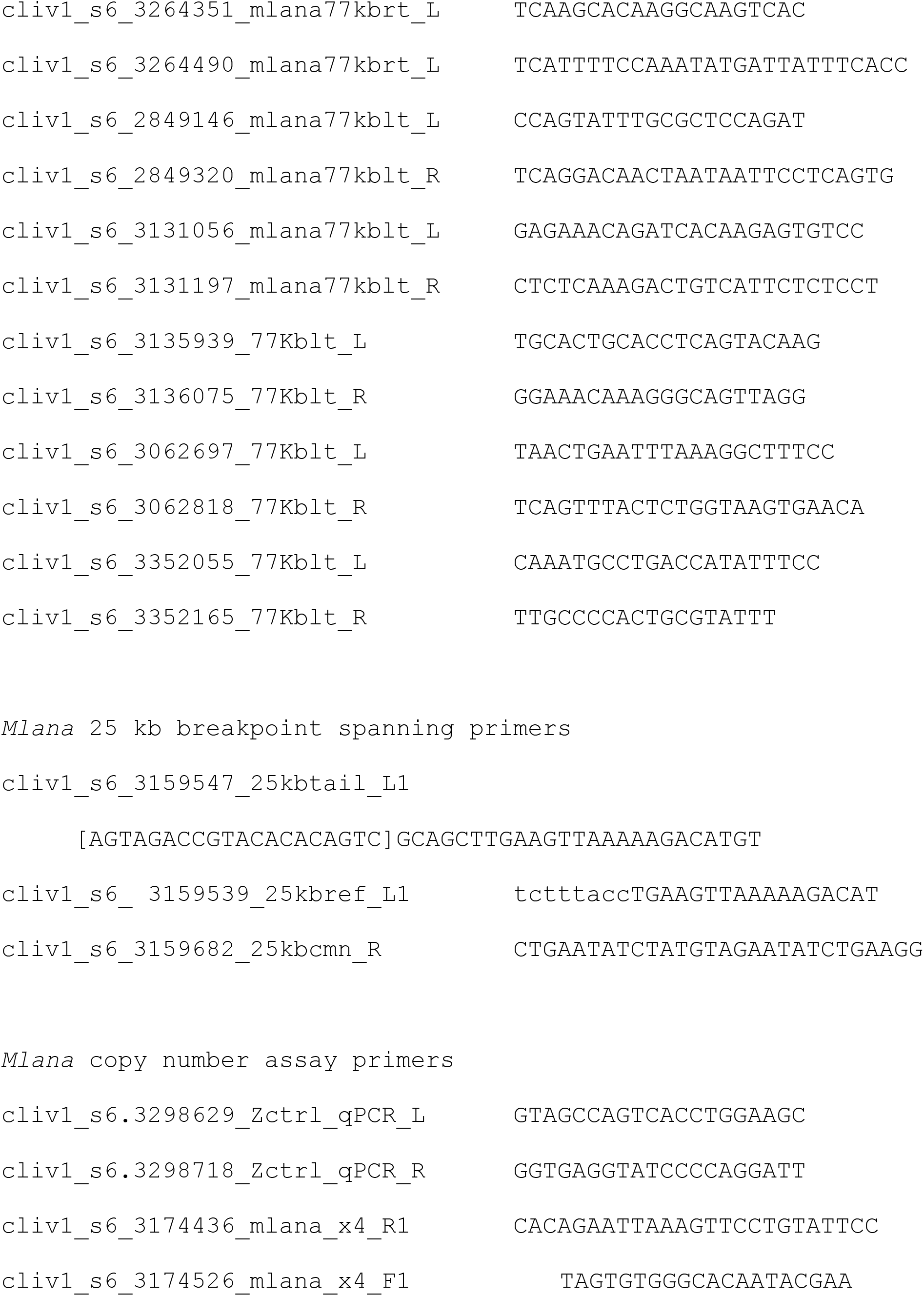

